# LANTHANUM (LaCl_3_) ADDITION DIVERSIFIES ORGANIC ACID PRODUCTION AND SIGNIFICANTLY ENHANCES METHANE PRODUCTION IN A METHANOGENIC CONSORTIUM

**DOI:** 10.64898/2026.08.24.746690

**Authors:** James Lawrence, Vincenzo Pelagalli, Gavin Collins, Piet N.L. Lens

## Abstract

Trace elements, such as iron, nickel, and cobalt are known to regulate methanogenic activity in anaerobic digestors used for waste valorisation, but the potential role of rare earth elements remains poorly understood. This study investigated the effects of lanthanum (La) supplementation on biogas production, methane generation, volatile fatty acid (VFA) formation, and carbohydrate utilisation in anaerobic digestion (AD). Biomethane potential (BMP) assays conducted under mesophilic conditions (37°C) using methanogenic sludge granules, and glucose as substrate, were supplemented with 0.1, 1, 10, and 100 mg/L lanthanum chloride (LaCl□). Biogas production and composition was monitored over a 96-h incubation, while sacrificial, batch bioreactors were used to evaluate temporal VFA and carbohydrate profiles. La supplementation significantly enhanced biogas and methane production in a concentration-dependent manner. The highest cumulative biogas yield (478.9 mL, corresponding to 179.5 mL biogas/g COD) and methane production (285.7 mL, corresponding to 107.1 mL CH_4_/g COD) were observed with 100 mg/L LaCl□, corresponding to increases of 88.7% and 186%, respectively, compared with La-free controls. CO_2_ production also increased with La concentration, whereas hydrogen production remained comparatively low. Acetic and butyric acids represented the dominant fermentation products (80-88% of total VFAs), but profiles of accumulated VFA in the bioreactors diversified with La addition, including showing caproate production, indicating changed biodegradation dynamics in the methanogenic microbiome. These findings demonstrate that lanthanum can stimulate anaerobic digestion performance and methane generation, highlighting the potential as a novel trace element additive to enhance biogas production. Research is now required to elucidate the underlying microbial and biochemical mechanisms, and establish optimal dosing strategies for large-scale applications.

**Graphical Abstract:** Representation of the experimental set-up, La dosing, and main observations.

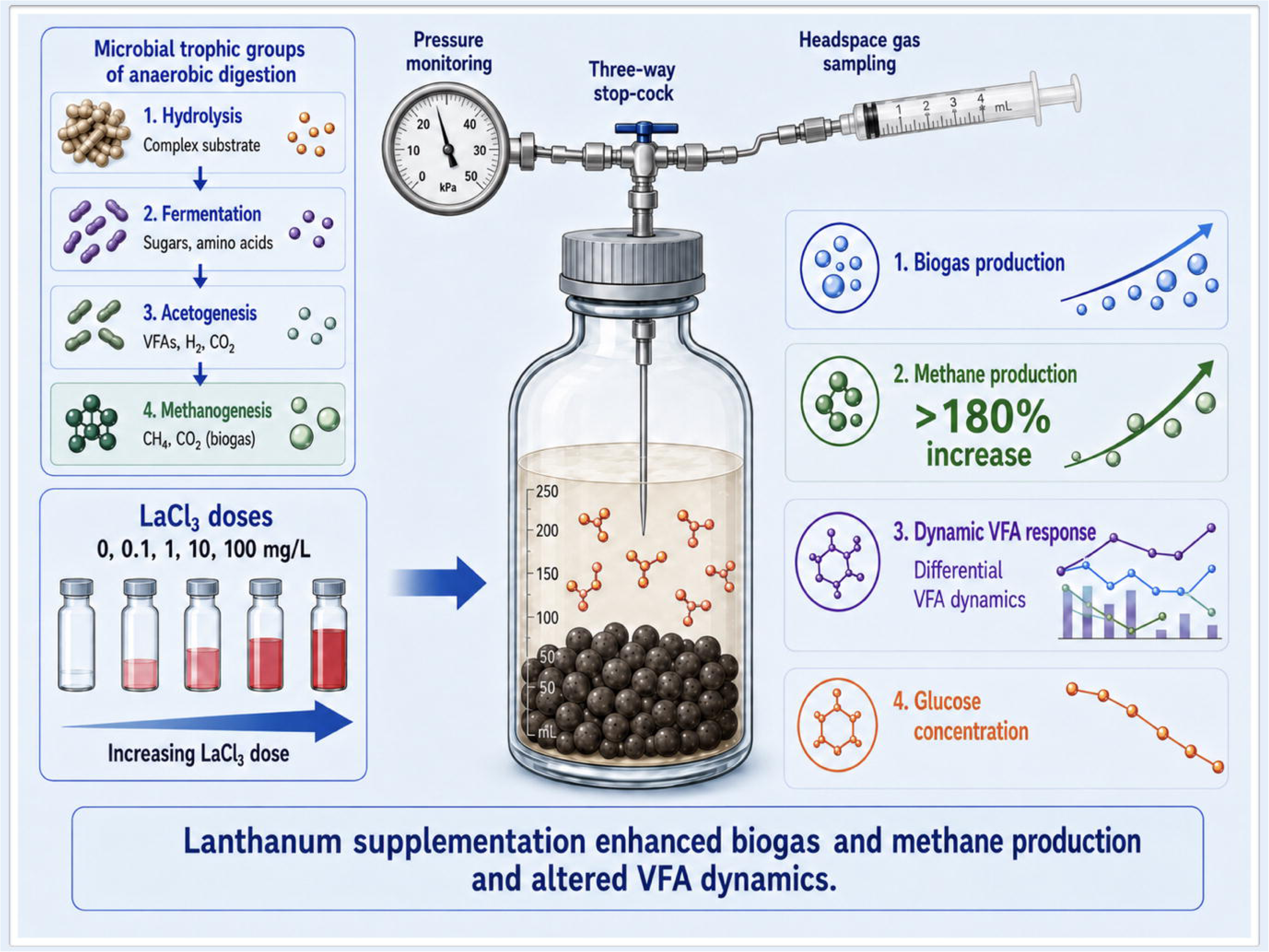

## 1. INTRODUCTION

Anaerobic digestion (AD) is a widely applied biotechnology converting organic polymers to biogas and digestate in the absence of oxygen and mediated by a series of microbial trophic groups, including hydrolytic, fermentative and acetogenic bacteria, and methanogenic archaea. Several challenges have impeded expanded application of AD technology, including conversion efficiencies and low methane (CH_4_) productivity, and process instability resulting from volatile fatty acids (VFAs) accumulation (Chang *et al*., 2025).

The presence in AD of various trace elements (TE), e.g., iron (Fe), nickel (Ni) and cobalt (CO), enhances methane production rates and yields, reduces lag times, and improves process stability (Song et al., 2025). Consequently, dosing of proprietary TE blends has emerged as key tool to optimise AD processes, providing cofactors for key metalloenzymes involved in electron transfer, redox balance, and carbon conversion pathways (Sun et al., 2023).

While the importance for methanogenic biology of transition metals such as Ni, Co, Fe, and molybdenum (Mo) has been extensively documented (Wintsche et al., 2016), the potential role of rare earth elements (REE) in AD remains comparatively underexplored. Such elements include the lanthanides (atomic numbers 57–71), scandium (Sc), and yttrium (Y), which have special chemical properties and are frequently used in industrial and agricultural settings (Shi et al., 2025). Lanthanum (La), a soft, silvery metal used in catalysts (Jadhav et al., 2025), has traditionally been considered biologically inert or even toxic at elevated concentrations (Malvandi et al., 2021). Recent findings have, however, changed this understanding, and shown La to be essential for aerobic methane oxidation (Nakagawa *et al.,* 2012; Pol *et al.,* 2014; Dandare et al., 2025). Lanthanides were shown to be crucial cofactors for pyrroloquinoline quinone (PQQ)-dependent methanol and alcohol dehydrogenases in methylotrophic bacteria, which provide greater catalytic efficiency than their calcium-dependent counterparts (Wehrmann et al., 2017). In addition, La, in the form of lanthanum oxide (La_2_O_3_) nanoparticles, can not only enrich electroactive syntrophic microbes but also act as a conductive bridge to facilitate the direct interspecies electron transfer (DIET) process between them (Yang et al., 2025). At present, there is growing evidence that REE may affect metabolic pathways and community structure in complex microbiomes, which could improve energy yields and substrate conversion (Yu and Chistoserdova, 2017; Dauman, 2019). However, the impact of REE on strictly anaerobic processes and biogas production has not thus far been systematically investigated.

In this study, the effects of La addition on the methanogenic activity of anaerobic granules and biogas production in an AD system were investigated. We investigated whether La functions as a stimulatory micronutrient, observing the impact on biogas and metabolic intermediaries. Our findings provide a fresh insight, inviting further research to investigate the potential for intensification of biogas generation and deeper understanding of REE microbiology.

## 2. MATERIALS AND METHODS

### 2.1 Experimental set-up

Anaerobic granular sludge collected from a dairy wastewater treatment plant operating at 37°C (Carbery Milk Products, Ballineen, Co.Cork) was used as inoculum. Glucose (Merck, Darmstadt, Germany) was used as organic substrate. Lanthanum chloride (LaCl_3_) (Merck, Darmstadt, Germany) was used to test the effect of increasing La concentrations on biomethanation of the substrate. An automated biomethane potential (BMP) system (Nautilus, Anaero Technology, UK) was used with compatible, 1-L vessels each with a working volume of 0.6 L as batch bioreactors. Each bioreactor was filled with 40 g inoculum (∼ 7 g VS L^-1^), 560 mL glucose solution (2.5 g L^-1^ or approx. 2.5 g COD L^-1^), and LaCl_3_ at one of five concentrations (0, 0.1, 1, 10, and 100 mg L^-1^).

The pH of the liquid was adjusted to 7.0 ± 0.2 by dosing 1M HCl or 3M NaOH. Bioreactors were closed with rubber stops and crimped with aluminium rings, and headspace was flushed with N_2_ gas. Five experimental replicates were set up for each of the La concentrations (*n* = 25). Incubations were at a constant temperature of 37°C as described by Logan et al. (2021).

Additional, sacrificially-sampled bioreactors, with the same conditions as described above (except not in the automated system), were set up to enable temporal sampling of bulk-liquid-phase samples at specific time points without altering the liquid:headspace volume, and to measure glucose and VFA concentrations. Samples were centrifuged at 12,000 rpm for 10 min. Supernatant was filtered using 0.22 µm-pore filters (Sarstedt, Nümbrecht, Germany) and frozen at -20°C.

### 2.2 Analytical methods

Biogas production was continuously monitored by the automatic system until the tests were concluded, as soon as biogas production reached a plateau (total duration of the tests approx. 96 h). Gas samples were taken periodically during the observation (2, 4, 6, 22, 26, 30, 44 h).

Gas samples were withdrawn through the sampling septum of the gas bags of the respective BMP assays using a gas-tight syringe, and immediately analysed to prevent leakage or compositional changes. Gas composition was determined using a 7890B gas chromatograph (Agilent, Santa Clara, USA) fitted with a thermal conductivity detector maintained at 250°C, allowing the quantification of CH□, CO□, N□, O□, and H□, following the method described by Braga and Lens (2023). Helium was employed as the carrier gas at a flow rate of 10 mL min□¹.

The pH of the bulk-liquid samples collected from each sacrificial bottle was determined using a pH-meter (pH/ORP 300, Cole-Parmer, Vernon Hills, USA) connected to a Hamilton SlimTrode electrode (Darmstadt, Germany).

Total carbohydrate measurements were made using the filtered samples to follow glucose consumption. The samples were mixed with a 5% phenol solution and sulfuric acid and subsequently incubated at 100°C following the Dubois method (Dubois et al., 1951). The carbohydrate content was determined by measuring the absorbance at 492 nm using a UV– VIS spectrophotometer UV-1900 (Shimadzu, Kyoto, Japan), with analysis done in triplicate as described by Lawrence *et al*. (2025).

VFA profiles were quantitatively determined in the filtered samples following the procedure described by Oliva *et al*. (2023). Analyses were performed using a 1260 Infinity II high-performance liquid chromatography (HPLC) system (Agilent, Santa Clara, USA) fitted with a Hi-Plex H column (300 × 7.7 mm). The column temperature was maintained at 60°C, and detection was carried out using a refractive index detector operated at 55°C. A 0.005 M H□SO□ solution served as mobile phase at a flow rate of 0.7 mL min ¹.

## 3. RESULTS

### 3.1 Impact of lanthanum dosing on gas production

Gas analysis indicated that hydrogen production was consistently the lowest among the gases measured (Fig. 1). CO□ was the most abundant gas detected after methane, and was elevated roughly in line with La dosing (Fig. 1).

**Figure 1.**
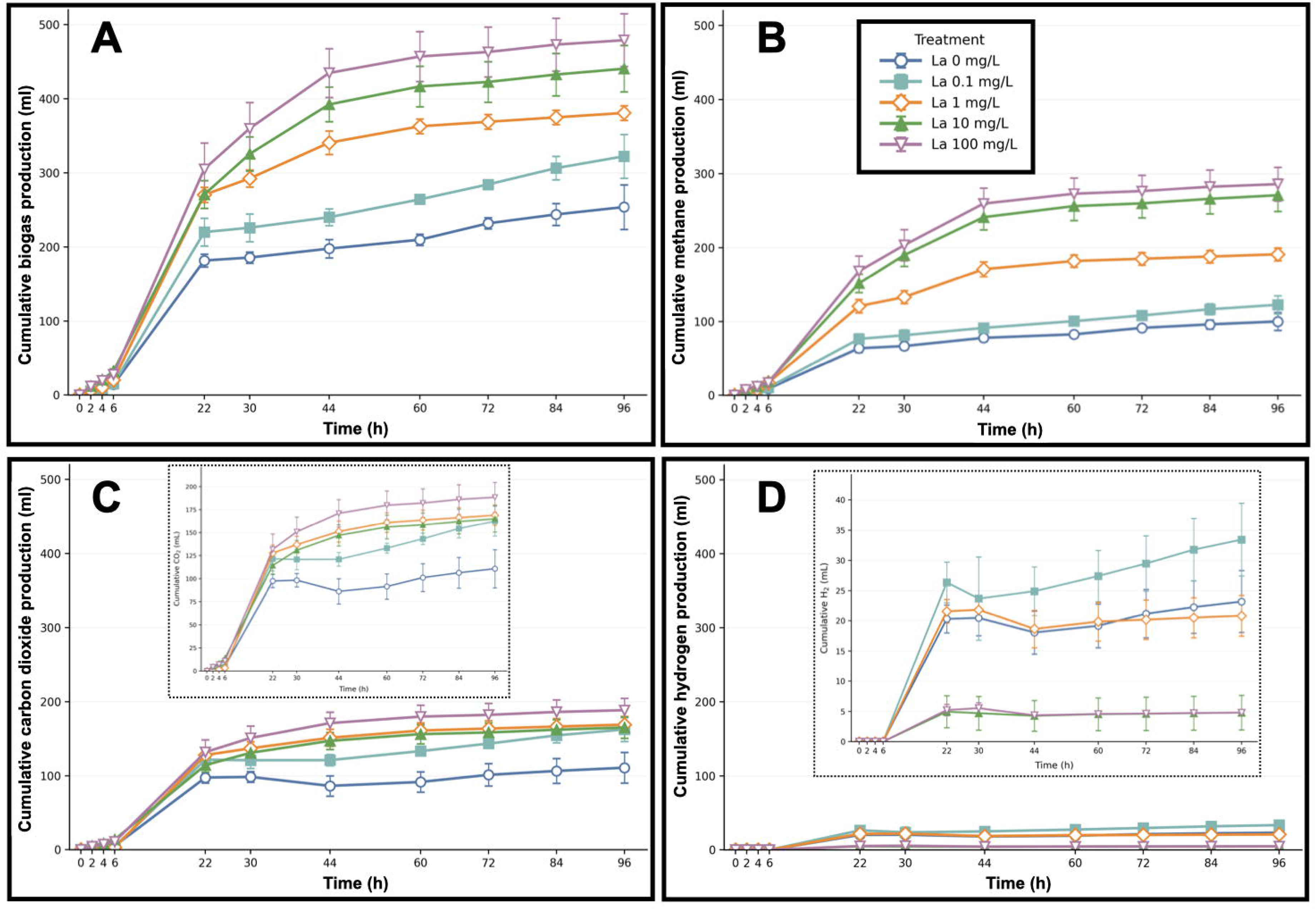
Cumulative biogas (A), methane (B), hydrogen (C) and carbon dioxide (D) curves obtained from La + glucose tests. Insets in panels C and D provide smaller y-axes to magnify the data from carbon dioxide and hydrogen production curves, respectively.

The addition of 100 mg/L LaCl_3_ resulted in production of the highest cumulative biogas volume (478.9 m/L or 179.5 mL biogas/g COD), followed by 10, 1 and 0.1 mg/L LaCl_3_, and the control (0 mg) (Fig. 1) – representing an 88.7% increase in gas production with supplementation at 100 mg/L LaCl_3_ compared to the La-free control. Specifically, methane production was also enhanced in response to La dosing. The highest methane production was observed with 100 mg/L LaCl_3_ (285.7 mL or 107.1 mL CH_4_/g COD), followed by 10, 1 and 0.1 mg/L LaCl_3_, and the control (0 mg) (Fig. 1) – representing a 186% increase in methane production compared with the La-free control.

Based on the standard theoretical maximum yield of methane from organic matter under anaerobic conditions at Standard Temperature and Pressure (STP) (Rittman and McCarty, 2001), the maximum production of methane from the COD supplied (2.5 g) was approximately 875 ml. The highest cumulative methane produced, which was with 100 mg/L LaCl_3_, was approx. 285 ml CH_4_, representing just above 30% methane yield efficiency.

### 3.2 Impact of lanthanum dosing on VFAs and glucose dynamics

Meanwhile, about 1.2 g of VFA-COD accumulated in the tests, representing a potential pool of over 400 ml CH_4_, accounting for most of the COD in the system.

The dominant VFA accumulated across the incubations with 0, 1 and 10 mg LaCl_3_ /L were butyric, acetic, propionic and iso-butyric acids (Fig. 2), and glucose was consumed under all conditions although more slowly in the presence of La. There was little difference in the concentration of total VFA accumulated in the tests with different LaCl_3_ applications, but LaCl_3_ dosing resulted in diversified VFA profiles characterised by accumulation of valeric, iso-valeric and caproic acids (Fig. 2).

**Figure 2.**
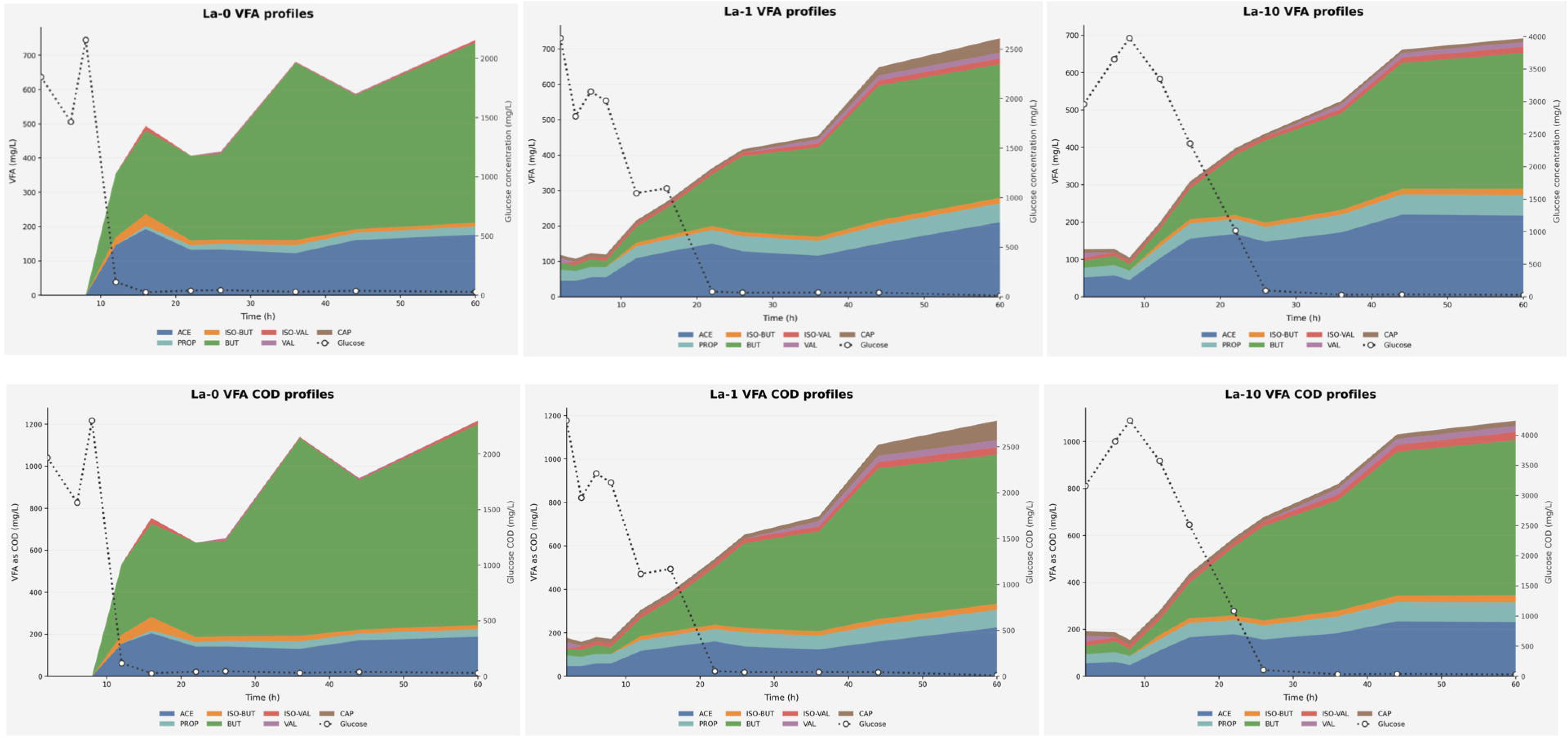
Temporal profiles of accumulated volatile fatty acids (VFAs) (A – C) and associated COD profiles (D – F) in La + glucose tests in the control (A, D), 1 mg/L (B, E) and 10 mg/L (C, F) La tests.

## 4. DISCUSSION

### 4.1 Effect of lanthanum addition on biogas and methane production

Interest in La as an additive in AD has grown in recent years, although the reported impacts are variable and require further interpretation. At low concentrations, lanthanum is frequently reported to stimulate methanogenic activity, with correspondingly increased gas yields and reduced lag phases (Yang *et al.,* 2022). Su *et al*. (2023) observed that supplementing low concentrations of La O and CeO (0–0.05 g/L) enhanced methanogenesis by approximately 4% and 3%, respectively. They proposed the observed effects were partly associated with the dissolution of La and Ce from the metal oxides, resulting in uptake and accumulation in the anaerobic sludge. Additionally, La O was more soluble and accumulated more in the anaerobic sludge. Similarly, in our study, LaCl may provide soluble La directly to microbial cells, influencing diversified VFA dynamics (Fig. 2) and elevated methane production (Fig. 1).

The stimulatory effects of lanthanides on methanogenic activity have been attributed to their ability to interact with redox-active enzymes, potentially enhancing key metabolic pathways involved in methanogenesis (Martinez-Gomez *et al.,* 2016). However, the beneficial window appears narrow. Beyond optimal concentrations, lanthanum may be inhibitory – for example, by precipitation with phosphate or disruption of trace nutrient balance, ultimately constraining microbial activity (Rajendran *et al.,* 2024). Similarly, Xing *et al*. (2022), indicated that 500 mg La (III)/L inhibited methanogenesis from cellulose, reducing methane production by 20%.

In a bioreactor study, Su *et al*. (2024) observed that La O addition resulted in an 11% reduction in COD removal and a 38% decrease in methane production, accompanied by disruption of extracellular polymeric substances (EPS) and sludge-granule (biofilm) disintegration. Despite this, they reported the La O induced potentially beneficial microbial changes, including an increased abundance of methanogens, suppression of propionate-and butyrate-producing bacteria, and stimulation of *Geobacter* and *Methanospirillum*. These contrasting findings suggest that lanthanum should not be considered a universally beneficial additive, but rather that performance depends strongly on system chemistry and dosing strategy.

The combined use of lanthanum with biochar introduces an additional layer of complexity, while also highlighting potentially synergistic effects. Biochar is well established as a conductive material that can promote direct interspecies electron transfer (DIET), thereby enhancing methanogenic efficiency (Zhang *et al.,* 2023). While few studies to date have focused the effects on methanogenic consortia of lanthanum-engineered biochar, a substantial body of literature has examined its adsorption properties (Wang *et al.,* 2016). Several potential benefits are associated with lanthanum modification, particularly with respect to enhanced contaminant removal and nutrient adsorption, which may have implications for anaerobic treatment systems. The strong affinity of lanthanum for oxyanions or Lewis bases has attracted significant attention in environmental applications, particularly for the adsorption of oxyanions or hard Lewis bases, such as phosphate, arsenic, antiomate, and fluoride (Yang *et al.,* 2023). Furthermore, the propensity of biochar to adsorb inhibitory compounds and buffer pH may alleviate potential toxicity associated with lanthanum accumulation (Zhao *et al.,* 2020; Balusamy *et al.,* 2015).

From the perspective of microbiome dynamics, lanthanum addition is associated with shifts in archaeal and bacterial communities. Xing *et al*. (2022) observed the enrichment of key acetoclastic methanogens (48.44%), such as *Methanosarcina*, *Methanothrix*, *Methanomassiliicoccus*, *Methanofollis*, *Methanobrevibacter*, *Methanobacterium*, and *Methanoculleus,* while the relative abundance of hydrogenotrophic archaea reached up to 45.35%, implying a stimulation of both acetoclastic and hydrogenotrophic methanogenesis pathways by La (III). Furthermore, lanthanum has also associated with elevated relative abundance of bacteria (*Acidobacteria* spp.) and fungal phyla in soil microbiomes (Song *et al.,* 2025). These changes are frequently more pronounced in the presence of biochar, supporting the idea that conductive materials strengthen microbial interactions through DIET (Gahlot *et al.,* 2020). Nevertheless, while lanthanum shows potential as a functional additive, its effects on biogas production and microbial structure are multifaceted and require further mechanistic investigation.

### 4.2 Impact of lanthanum on volatile fatty acid conversion

As a trivalent rare earth element, lanthanum can interact with microbial enzymatic systems by substituting for essential metal cofactors such as calcium due to its comparable ionic radius (Nikolova *et al.,* 2023). This substitution can alter enzyme conformation and catalytic efficiency, thereby affecting key biochemical pathways. Notably, lanthanides have been shown to enhance the activity of specific dehydrogenases, especially those involved in oxidation–reduction reactions, which play a central role in VFA formation.

Ji *et al*. (2024) demonstrated that moderate nickel supplementation with iron-doped lanthanum manganese oxide nanoparticles can promote pyruvic acid production from glucose with conversion to VFA and H_2_, finding that appropriate dosage of LaMn_0.7_Fe_0.3_O_3_ could significantly improve glucose metabolism.

### 4.3 Comparison with conventional trace element supplementation

The potential of La in supplementation of methanogenic consortia appears highly dependent on organic substrate type and bioreactor conditions (Xing *et al.,* 2022). For example, improved carbohydrate degradation and increased acetate formation have been linked to lanthanide-dependent enzymatic activity, which can feed into methanogenic pathways (Ji *et al.,* 2024). Traditional trace elements serve as well-established co-factors in enzymatic processes, such as methanogenesis, but lanthanum appears associated with more selective control over microbial pathways, particularly those involving redox-active enzymes (Martinez-Gomez *et al.,* 2016). Comparative studies suggest that, at optimized concentrations (below 500 mg/L), lanthanum supplementation can enhance methane production and overall gas yields relative to conventional trace elements (Xing *et al.,* 2022).

Unlike iron, cobalt, and nickel which have well-defined biological roles in enzymes such as hydrogenases and methyl-coenzyme M reductase, the function of lanthanum is less clearly established and more variable (Myszograj *et al.,* 2018). The effects of La are strongly concentration-dependent, where low levels may stimulate microbial growth and enzymatic efficiency enhancing alternative pathways such as acetate decarboxylation and utilization of H_2_, methanol, and methylamine, while higher concentrations can inhibit methanogenic archaea and gas production (Yang *et al.,* 2026; Crombie *et al.,* 2022). This variability underscores the importance of precise dosing in trace element management strategies.

Among the studies reviewed here (Table 1), Chan *et al*. (2019) reported one of the most substantial improvements in gas production, observing 94.1% increase following the addition of copper (10 mg/L) to domestic food waste and activated sludge. Similarly, Lizama *et al*. (2025) reported 75.8% enhancement in biogas yield with cobalt supplementation (7 mg/g VS) in sewage sludge. Although the absolute biogas yield obtained in the present study was lower than those reported in those investigations, methane production increased by 186% relative to the control treatment, indicating a pronounced stimulatory effect of lanthanum supplementation under the conditions tested.

**Table 1.** Comparative improvements of biogas yields across trace element supplementations of methanogenic systems.

| Organic substrate | Inoculum source | Trace element supplied | Biogas yield | Yield increase | Reference |
| --- | --- | --- | --- | --- | --- |
| Sewage sludge | Waste treatment plant sludge | Iron (Fe) – Fe<sup>0</sup><br>10 g/L | 165.1 mL/g VSS | **13.2%** | Zhang et al., 2014 |
| Straw | Cow dung | Nickel (Ni) -<br>NiCl<sub>2</sub> 2 mg<br>Ni/L | 32.44 mL/g TS | **18%** | Tian et al., 2017 |
| Food waste | Digestate | Selenium (Se)<br>–10 – 20 µg/L<br>(0.13 – 0.25 µM) | 537 mL/g VS | **30.1 %** | Lens et al., 2016 |
| Sewage sludge | Sewage sludge | Cobalt (Co) - 7<br>mg/g VS | 232 mL/g VS | **75.8%** | Lizama et al., 2025 |
| Food waste and domestic wastewater | Activated sludge | Copper (Cu) -<br>Cu<sup>2+</sup> 10 mg/L | 260–325 mL<br>CH<sub>4</sub>/g COD | **94.1%** | Chan et al., 2019 |
| Food waste | Seed sludge | Molybdenum (Mo) – 5 mg/L | 415 ± 5 mL/g VS | **11.6 %** | Zhang et al., 2015 |
| Food waste | Seed sludge | Trace element mix<sup>a</sup> | 504 mL/g VS | **35.5%** | Zhang et al., 2015 |
| Food waste | Commercial AD plant sludge | Trace element mix<sup>b</sup> | 677.9 mL/g VS | **27.4 %** | Zhu et al., 2024 |
| Synthetic wastewater | Anaerobic granular sludge | Lanthanum -<br>La<sub>2</sub>O<sub>3</sub> | 56.26<br>mL/(h·gVSS) | **4%** | Su et al., 2023 |
| Glucose | Anaerobic granules | Lanthanum –<br>LaCl<sub>3</sub> 100<br>mg/L | 107.1 mL CH<sub>4</sub>/g<br>COD | **186%** | This study |

<sup>a</sup> Fe (100 mg/L) + Co (1 mg/L) + Mo (5 mg/L) + Ni (5 mg/L).
<sup>b</sup> Fe (113.1 mg/L) + Co (0.51 mg/L) + Ni (2.44 mg/L).

From a management perspective, incorporating lanthanum into supplementation regimes presents both opportunities and challenges. Insights from metatranscriptomic analyses further reveal that lanthanum can regulate microbial metabolism. For example, lanthanide availability was shown to upregulate genes associated with alcohol dehydrogenases and other key enzymes involved in carbon metabolism, while downregulating less efficient metabolic routes (Gorniak et al., 2023). These shifts suggest lanthanum not only affects enzymatic activity but also reshapes metabolic networks, contributing to improved substrate conversion and gas production under optimal conditions. Indeed, La has been reported to stabilize enzyme structures, improve catalytic activity, and enhance microbial growth (Hu et al., 2017). It may also influence the balance of reactive oxygen species (ROS) within microbial systems (Liu et al., 2016). Despite low oxygen concentrations under anoxic conditions, oxidative stress can still occur due to metabolic activity or environmental fluctuations (Fu et al., 2015). Evidence suggests that low concentrations of lanthanum can enhance antioxidant defences, possibly by stimulating enzymes such as superoxide dismutase, thereby protecting cells from ROS damage (Liu et al., 2016). In contrast, excessive lanthanum may induce oxidative stress by interfering with cellular metal homeostasis, leading to impaired metabolic function (Todorov et al., 2019). Furthermore, the introduction of lanthanum may allow for reduced reliance on conventional trace metals or enable targeted manipulation of metabolic pathways to improve process efficiency. However, potential interactions with existing trace elements must also be carefully considered, as lanthanum can compete with essential cations, such as calcium and magnesium, potentially disrupting enzymatic balance and microbial homeostasis (Niklova et al., 2023). Therefore, system-specific optimization and monitoring will be critical for successful implementation.

Studies aimed at optimizing the fundamental parameters governing microbial growth and enrichment conditions are limited. To date, available evidence has primarily focused on the roles of trace metals, particularly iron, copper, and lanthanides, in influencing microbial physiology and metabolic activity (Hatamoto et al., 2018; Guerrero-Cruz et al., 2021). However, prospective dosing will require a thorough understanding of concentration-dependent effects, interactions with other micronutrients, and long-term impacts on microbiome dynamics and function. Future research integrating biochemical, physiological, ‘omics, and modelling will be essential to fully harness the potential of La dosing in AD systems.

## CONCLUSION

Lanthanum presents promising prospects as a supplementary or alternative trace element in anaerobic digestion. Lanthanum supplementation enhanced the activity of a methanogenic consortium, resulting in significantly enhanced biogas and methane production, and altered fermentation dynamics. The highest La concentration tested (100 mg/L) enhanced methane production by 186%, indicating La may act as a stimulatory trace element in methanogenic systems. The observations highlight the potential of lanthanum as a novel additive for improved biogas production, although deeper studies are needed to elucidate the underlying mechanisms and determine optimal application strategies.

## Acknowledgements

Mr Borja Khatabi Soliman is thanked for his assistance in executing the incubations used in this experiment.

## Author contributions

PL and GC conceived of the experimental question and design. VP and JL executed the experimental work. JL drafted the manuscript, and all authors contributed to data analysis and manuscript editing. PL (15/RP/2763 and 2023GEH252) and GC (25/FIP/GNI/15297) were responsible for funding acquisition.

## Data availability

The data underpinning this article are available upon request.

